# Towards Fully Automated Investigation of Social Learning in Mice

**DOI:** 10.1101/2025.08.31.673387

**Authors:** Benjamin Lang, Christa Thöne-Reineke, Olaf Hellwich, Lars Lewejohann

**Affiliations:** Science of Intelligence, Research Cluster of Excellence, Marchstr. 23, 10587 Berlin, Germany; Institute of Software Engineering and Theoretical Computer Science, Faculty IV – Software Engineering and Theoretical Computer Science, Technische Universität Berlin, Berlin, Germany; Institute of Animal Welfare, Animal Behavior and Laboratory Animal Science, School of Veterinary Medicine, Freie Universität Berlin, Berlin, Germany; Institute of Computer Engineering and Microelectronics, Faculty IV – Software Engineering and Theoretical Computer Science, Technische Universität Berlin, Berlin, Germany; German Centre for the Protection of Laboratory Animals (Bf3R), German Federal Institute for Risk Assessment (BfR), Berlin, Germany

**Keywords:** Social Learning, Mice, Home-cage monitoring, Animal Tracking, IntelliCage, Live Mouse Tracker

## Abstract

Mice have been demonstrated to learn from each other in social interactions, the extent to which this takes place and the strategies involved, however, largely remain to be elucidated beyond spatially and temporally confined tests of social memory retention. Here, we present a method which utilizes and modifies 1) a commercially available tool for automated behavioral testing, the IntelliCage and 2) an open-source solution for 24/7 live animal tracking, the Live Mouse Tracker, to create a powerful method for the investigation of learning behavior in semi-naturalistic group settings. We see a wide range of possible applications, such as for instance the investigation of learning in social interactions, which we present here as an example. In the present study, co-learning did not facilitate place learning over individual learning. While automated annotation of behaviors was effective, markerless animal identification proved unreliable in a highly enriched environment with manifold opportunity of occlusion from video tracking. In response, we here present a rationale for identifying the reliable portion of tracking data, to which we confine the behavioral analysis. Co-learning animals engaged more often in some prosocial interactions with their teammates than with other animals, but they did overall not interact with each other more frequently than with individual learners. Correlative analysis of learning behavior and social interactions did not reveal any particular association between behavior and learning success. While the mechanisms of social learning in mice could not be conclusively elucidated within the scope of this study, we report on the development of a promising tool for presenting manifold learning tasks to mice while tracking their individual and social behaviors in a fully automated manner. Further, we discuss limitations of the current configuration and present an outlook on further improving the method.

## 1 Introduction

### Social Learning in Mice

Mice (*Mus musculus*) are by far the most often used mammalian species in research(1). Their relative genetic similarity to humans (2), short generation times and comparatively low demands on housing and maintenance (3) make them a fitting choice for manifold research questions in biomedical research, compound testing, and biological- as well as psychological basic research. In the wild, mice live in large social groups (4) with complex hierarchies (5,6). The evolution of their social lifestyle entailed mechanisms for the social transmission of information and learning from social interactions with conspecifics. Those in turn are utilized in biomedical research to test for impairments of social behavior associated with e.g., novel compounds (7) or transgenic models for autism (8). Most prevalent are tests of social amnesia in dyadic or triadic interactions in a designated experimental area distinct from the animals’ home cage, thus being spatially and temporally confined (9). Neurologically unimpaired rodents generally display greater interest in unfamiliar conspecifics than in familiar ones. In such assays, as for instance the ‘three-chambered social memory test’ (10), decreased social memory retention generally manifests as prolonged exploratory behavior towards familiar animals, which is used to analyze impairments of social behavior.

While the utility of such assays for screening for impaired social memory is undisputed, they are limited in the scope of behaviors they can probe. In recent years, advancements in automation and animal tracking have furthered efforts to address the limitations of ‘classical’ dyadic social tests by offering a more comprehensive insight into mouse behaviors. Examples for animal tracking solutions seeing widespread use include DeepLabCut (11), LEAP (12), and SuperAnimal (13).

The extent to which mice are capable of inferring information from their conspecifics remains inconclusive. Mice have been demonstrated to display preference towards food in choice tests for items that were previously consumed by conspecifics (14) and to socially learn fear of adverse stimuli (15). Learning of more complex social behaviors is sparsely documented. Few studies found decreased latency in solving baited mechanical puzzles in mice previously given the chance to observe a demonstrator conspecific as compared to naïve mice (14,16,17). None of those studies however observed true copying of behavior in the mice, and they all probed social learning in controlled interactions. Studies which investigate social learning in more semi-naturalistic group settings are even more sparse (18). However, with the increasing availability of multi-animal tracking solutions and methods for automated behavioral testing, we anticipate an increase in studies of socially-housed groups of mice offering a more comprehensive understanding of murine cognition and behavior (reviewed in (19)).

### IntelliCage

The IntelliCage (IC) is a home-cage-based system for automated behavioral testing of rodents. It is commercially available through TSE Systems (20). It allows for the fully automated presentation of place learning tasks as well as automated acquisition of data, thereby increasing reproducibility by minimizing experimenter interference (21). The IC consists of a controller unit and four ‘operant corners’, embedded in an aluminum cover. The entire unit can be placed in a Makrolon type IV cage and house social groups of up to 16 animals. Each operant corner contains two bottles which can be filled with liquid rewards (e.g., H_2_O, saccharose solution, almond milk), which can be accessed through a retractable door each. The doors can be programmed to open for individual animals or groups once certain conditions are met. Each corner holds a presence sensor as well as a radio frequency identification (RFID) antenna for animal identification, a light-barrier sensor for detection of ‘nose pokes’ (NPs) by the animals, a so-called ‘lickometer’ for the monitoring of animal liquid intake, and three colored LEDs for giving light signals. Further, aversive stimuli can be delivered via pressurized air (air puffs). The IC can be programmed to give access to the liquid rewards e.g., at a certain location, to a certain time, once a certain number of NPs is performed, or in more complex shuffle patterns; thereby allowing for spatial- and temporal learning paradigms. Errors can be punished by withholding reward or delivering air puffs. One limitation of the IC is that all sensors are confined to the operant corners, not offering any way to monitor behavior and social interactions taking place in the main living space of the IC.

### Live Mouse Tracker

The Live Mouse Tracker (LMT) was developed by de Chaumont *et al*. (22) as a method for 24/7 live tracking and automated behavioral annotation of groups of mice. The method is described in detail in the complementary publication (22), as well as on Fabrice de Chaumont’s website (23). Briefly, it uses a depth-sensing infrared camera supported by computer vision and machine learning in conjunction with 16 RFID antennae for 24/7 identification, tracking and automated behavioral annotation of mice. In the current setup, it can track 4 mice at a time and is constructed to be used with a custom-built 50 × 50 cm cage. Since its introduction in 2019, it has for instance been used to investigate the contributions of autism-associated mutations in the *Shank2* and *Shank3* genes (22) as well as exposure to valproic acid to social behavior deficits (24).

Here, we report on a method which builds on both the IC and LMT to achieve fully automated behavioral testing with continuous multi-animal tracking. We use this method to test whether co-learning scenarios in mice facilitate decision making and development of a strategy to exploit dynamic probabilistic reward opportunities. We evaluate the method’s performance in a highly enriched environment and identify caveats, which lead us to only exemplarily evaluate animal behavior. Yet, we are confident to present a powerful tool with manifold potential applications reaching from investigations of social impairments to testing models of social information transmission.

## 2 Methods

### Ethics Statement

All experiments involving animals were carried out in accordance with the German Animal Welfare Act and Directive 2010/63/EU. Approval for animal maintenance and experimentation was granted by the Berlin state authority (Landesamt für Gesundheit und Soziales) under permit number 0249/19. Prior to the start of the experiments, the study was pre-registered at animalstudyregistry.org (DOI: 10.17590/asr.0000283).

### Animals

16 female pathogen-free C57BL/6J mice were purchased at the age of 42 - 48 d from Charles River Deutschland GmbH, Sulzfeld. The experiments were conducted with two batches (B1 and B2) of 8 mice, which underwent testing successively. Upon arrival, the animals of each batch were randomly divided into two groups (n = 4; groups 1 and 2). In accordance with this nomenclature, the 4 social groups are herein referred to as B1.1, B1.2, B2.1 and B2.2. The animals were handled via a 11 × 4 cm polymethyl methacrylate (PMMA) tube and trained to enter the tube voluntarily during the first two weeks after arrival.

### Animal Housing and Enrichment

#### Housing

Animals were kept in an animal facility under pathogen-free barrier conditions at 22 ± 1°C and 55 ± 1 % humidity. They were subjected to a 12 h light/dark cycle (light phase from 07:00 – 19:00). In the 30 min leading up to the beginning of the light phase, sunrise was simulated by a wake-up light (Philips HF 3510, 100–240 vac, 50–60 Hz, Philips GmbH Market DACH, Hamburg, Germany). During the acclimatization phase, animals were housed in a Makrolon type III cage (39 × 23 × 15 cm) with food (LASvendi, LAS QCDiet, Rod 16, autoclavable) and tap water *ad libitum*. At the age of 68 – 74 d, they were moved to the LMT-IC experimental setup where they lived permanently for the duration of the study. Animals were only briefly removed from the experimental apparatus for weekly cleaning and health checks. During the experimental phase, food (LASvendi, LAS QCDiet, Rod 16, autoclavable) was available *ad libitum* and tap water/4 % saccharose in H_2_O was available via the reward bottles of the IC.

### Enrichment

**Fig. 1C** depicts the 49.2 × 49.2 cm enriched living area. The living area was filled with 5 l of fine wooden bedding (JRS Lignocel FS14, spruce/fir, 2.5 – 4 mm). A main nesting area was provided consisting of three houses (1× Fat Rat Hut; 16 × 8.6 × 15.2 cm; Bio-Serv, Flemington, New Jersey, USA; 2× triangular Mouse House; 14.5/11.5 × 5 cm; Tecniplast) containing three thin paper towels (cellulose, unbleached, 20 × 20 cm; Lohmann & Rauscher, Neuwied, Germany). In two orthogonal corners of the living area, two additional smaller houses (Safe Harbor Mouse Retreat; 6.4 × 11.4 × 7 cm; Bio-Serv, Flemington, New Jersey, USA) were provided, further a mouse house/running disk combination (house: 10.5 × 5.5 cm, disk: 15 cm; ZOONLAB GmbH, Castrop-Rauxel, Germany). Per cage, 8 wooden gnawing blocks were supplied as well as additional nesting material in the form of 12 cotton rolls (Gr.3, UNIGLOVES, Troisdorf, Germany) and 30 paper strips (LILLICO, Biotechnology Paper Wool). Further, the PMMA tube used for animal handling remained in the cage. At two time points (d1 and d21 of the co-learning experiment), a 3D-printed puzzle ball from which the animals could retrieve millet (Alnatura GmbH, Darmstadt, Germany) was offered as additional enrichment.

**Fig. 1.**
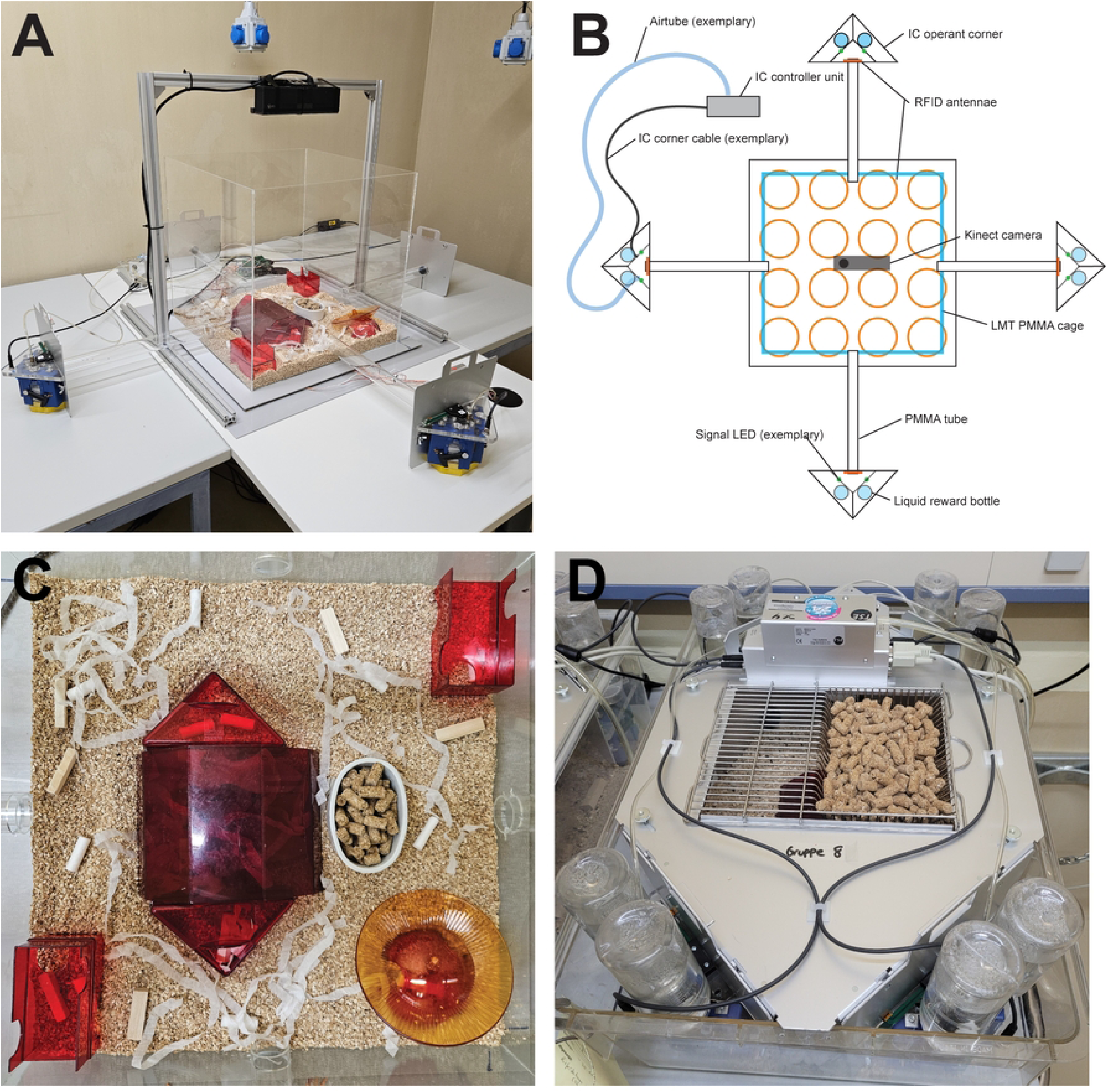
LMT-IC Apparatus. (A) Overview of LMT-IC setup with enrichment. (B) A simplified schematic of the LMT-IC setup from the top-down view. (C) Enrichment of the LMT-IC setup. Food was offered in a bowl on the bottom rather than in a rack to prevent occlusion from the camera. (D) External view of an IC in unmodified original configuration.

### Animal Identification

After one week of acclimatation to the animal facility (at 49 - 56 d of age), animals were subcutaneously implanted in the dorsocervical region with radiofrequency identification (RFID) transponders through application cannulas (2.6 × 0.15 × 40 mm ISO-Compliant Transponder; Peddymark Limited, Redhill, Surrey, United Kingdom).

Implantation was carried out under isoflurane narcosis (induction: 4 % isoflurane CP 1ml/ml in oxygen at 2 l/min; maintenance: 1 % isoflurane CP 1ml/ml in oxygen at 1 l/min; CP-Pharma Handelsgesellschaft GmbH, Burgdorf, Germany). ∼12 h before implantation, animals were orally treated with 0.5 mgkg^-1^ Metacam 0.5 mg/ml Suspension (Boehringer Ingelheim, Ingelheim am Rhein, Germany). Additionally, animals were marked with colors (red, green, blue or yellow; edding 750 paint marker, edding International GmbH, Ahrensburg, Germany) for quick visual identification by experimenters and animal caretakers at the tail base. Markings were maintained weekly.

### Live Mouse Tracker

Two Live Mouse Tracker (LMT) apparatuses were constructed as described in detail by Chaumont *et al*. (25). Briefly, per apparatus, a custom cage with a living area of 49.2 × 49.2 cm was constructed out of transparent Ø4 mm PMMA (Firstlaser GmbH, Bardowick, Hamburg, Germany). Located below the cage was an array of 16 (4 × 4) 100 mm RFID antennae (Priority 1 Design, Port Melbourne, Melbourne, Australia) manually tuned to 134.2 Hz. Placed on a custom-built aluminum rack (Aluminum Profile 30 × 30L B-Type Groove 8, DOLD Mechatronik GmbH; Haslach, Germany) was a depth-sensing infrared camera (Xbox One Kinect Sensor 2.0, Microsoft Cooperation, Redmond, USA) with one of the infrared LEDs masked to avoid overexposure of the recordings. The system was controlled via the Live Mouse Tracker application (Version 1040). The original design by de Chaumont *et al*. was modified in the following ways: 1) the height of the PMMA side panels was increased by 10 cm (from 35 cm to 45 cm), and 2) a circular hole with radius = 2 cm was laser-cut centrally in all 4 side panels at 4 cm height. Via those holes, 40 × 4 cm PMMA tubes were connected to the IC operant corners.

### IntelliCage

ICs were purchased from TSE Systems (20). Two V1.4 models were used in the present study. Experiments were controlled by the complimentary IC Software V3.3.5.0.

### Experimental Setup

A method was devised to combine the capacity for fully automated testing of spatial memory offered by the IC with the 24/7 live tracking of the Live Mouse Tracker (herein, LMT-IC). To achieve this, the simple approach of making the IC operant corners accessible from the 49.2 x 49.2 cm living area of the LMT was chosen. Both systems (IC and LMT) rely on RFID for animal identification. The LMT uses an array of 16 100 mm antennae located below the bottom of the living area, which are sequentially activated in order to validate or correct, respectively, the identity of individual animals. The IC includes one RFID antenna at the entrance of each operant corner, which are active constantly during operation. Due to those designs, the RFID antennae of the LMT and the IC are oriented perpendicular to each other. The same is true for the electromagnetic fields projected by the antennae, which may result in interference of the fields (26), preventing correct reading of transponders within overlapping fields. Preliminary testing revealed the area of interference to be restricted to a distance of <40 cm. Hence, rather than placing the IC operant corners inside the living area of the LMT apparatus in a similar configuration as in the IC setup (**Fig. 1D**), we opted to place them outside the cage, in 40 cm distance to the outer edge of the living area. The operant corners were connected via PMMA tubes (42 × 4 cm, custom-built from GEHR PMMA XT® ACRYL, Mannheim, Germany), preventing interference of the constantly active IC RFID antennae with the RFID array of the LMT and allowing for reliable reading of animal transponders.

#### 3D printing

Custom sockets for external placement of the IC operant corners were designed (**Fig. 2**) in tinkerCAD (27) and 3D-printed (Ultimaker^3^ Extended; Freeform4U GmbH, Munich, Germany; filament: Polylactic acid (PLA); Filamentworld, Neu-Ulm, Germany). The sockets were designed in a way to ensure stability of the IC operant corners via pins intruding the door path of each operant corner. Inside those pins, there was an internal canal to allow for any excess liquid spilled during liquid consumption of the mice to flow down the canal and be collected in a reservoir at the bottom to prevent accumulation of liquid in the apparatus for hygienic reasons.

**Fig. 2.**
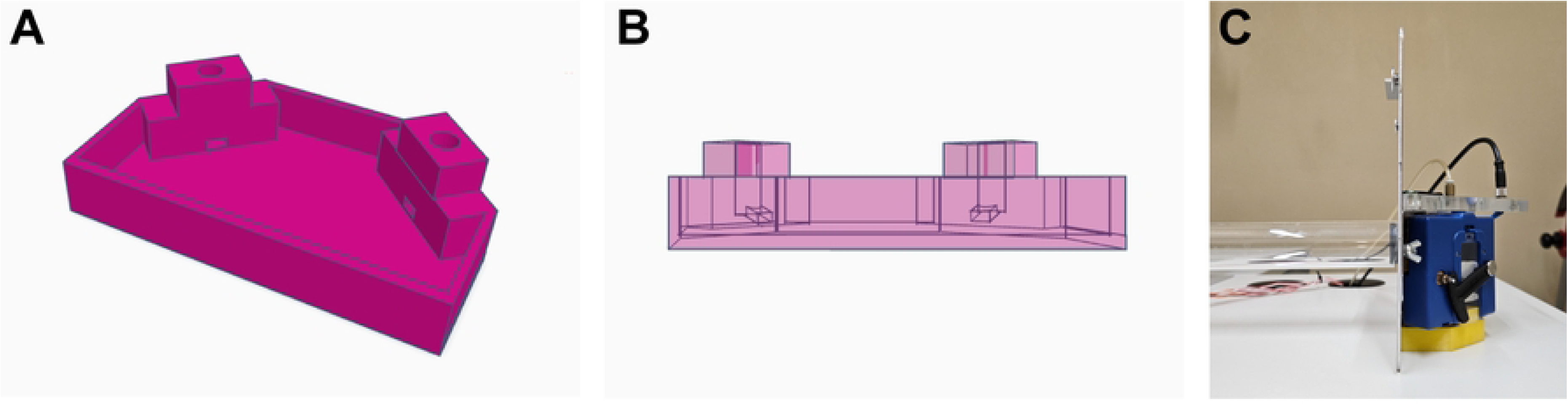
Custom socket for placement of IC operant corners outside of cage area. Custom sockets were designed and 3D-printed to allow connecting the IC operant corners externally to the cage area without the default IC aluminum frame. An internal canal leading into a reservoir allowed for the removal of any excess liquid. (A) 3D model of socket. (B) Translucent 3D model of socket with internal canal visible (C) IC operant corner placed in 3D-printed socket (yellow), connected to PMMA tube leading into the living area.

#### Custom modifications to the IC

The IC operant corners were removed from the aluminum cover and placed in the 3D- printed sockets. The controller unit was unscrewed from the aluminum cover for independent usage. All modifications done to the IC were reversible so that the IC could be used in its original configuration in other experiments. To accommodate for the greater dimensions of the LMT-based setup, longer variants of the cables connecting the IC main controller unit with the operant corners were soldered and custom tubes (for delivering airpuffs to the IC operant corners) longer than the original versions were deployed.

### Experimental procedure

The co-learning paradigm was carried out for 41 consecutive days and nights, during which 16 C57BL/6J mice were continuously living in the LMT-IC apparatus in groups of n= 4, each. Two groups at a time underwent testing simultaneously. Each IC operant corner held two liquid reward bottles, filled with 250 ml of 4 % saccharose in H_2_O. The reward bottles were accessible to the animals during 2 drinking phases each night reaching from 21:00 – 23:00 and 03:00 – 05:00, respectively. Prior to the co-learning paradigm, animals were gradually habituated to the experimental apparatus, to the drinking phases, to frequenting the operant corners as well as performing nose pokes (NP) in order to receive reward. Additionally, individual performance in the probabilistic place-learning task was recorded for w as baseline measurement before subjecting the animals to the co-learning paradigm.

At the start of the co-learning paradigm, animals were randomly assigned to either ‘team learning’ or ‘solo learning’. ‘Team learning’ here describes a co-learning paradigm in which two animals per group had the highest probability (p = 0.9) for receiving a reward at the same location. The remaining three operant corners all gave off reward with p = 0.1, each. The other two animals per group had the highest chance for reward at an individual corner each, not coinciding with the highest reward probability option of any other animal, acting as an in-cage control. Every third night, the distribution of reward probability options shifted across the operant corners in a semi-randomized manner: a new corner was assigned randomly, with the exclusion of the previously assigned corner and any high-reward opportunity operant corner assigned to another solo-learning animal. Low reward probability status was indicated to the animals upon entry of a corner with a green LED signal for 1 s. Doors to the reward bottles opened after performing 6 NPs to the light barrier sensor, to which the animals were gradually habituated prior to the start of the experiment. Doors closed after 6 s and would open again only after an animal left the operant corner and performed a new trial. Over the entire experimental period, animal movement and behavior were tracked via the depth-sensing infrared camera and RFID antennae of the LMT apparatus.

#### Rewards and aversive stimuli

During the acclimatization phase, all IC operant corners were baited with plain H_2_O. During baseline measurements and the co-learning paradigm, corners were baited with 4 % saccharose (Millipore Ltd Oakville, Ontario, USA) in H_2_O. The rationale behind offering saccharose was to 1) increase animal motivation to participate in the trial, 2) putatively give the animals additional social cues via saccharose olfaction (28) and increased affective signaling upon receiving saccharose reward. The 4 % saccharose solution was exchanged thrice per week and then freshly prepared by stirring it until it was completely dissolved. Per operant corner, 2 bottles with 250 ml of saccharose solution were deployed. Liquid intake was monitored daily. When an animal stayed at the same operant corner without leaving for >40 s, an airpfuff (1 s at 0.5 bar) was delivered to the respective operant corner. This was to discourage nesting behavior in the operant corners. During the habituation phase, animals were habituated to the 40 s window by gradually decreasing the time until the airpuff was delivered from 180 s to 40 s.

### Data Processing

#### IC Data

Learning data collected through the IC was analyzed in the open-source statistical software RStudio (Version 2024.04.2 Build 764). For graphical visualization of data, the ggplot2 (29) package was deployed. For statistical modelling, the mixed-effects model package lmerTest (30)was used. Model assumptions were checked visually by Q-Q-plotting and assessing variance homogeneity of residuals versus fitted values. The model assumed the nightly average preference for the high-reward probability option as response variable, with the fixed factors 1) social learning configuration (2 levels), 2) night number relative to the reward probability distribution changing (3 levels), absolute night number (32 levels), social group (4 levels), and the individual animals (n = 16) as random effect. The overall effects of each fixed factor were evaluated via Type-II F-tests with Satterthwaite degrees of freedom (30). For factors with significant effect, estimated marginal means were computed and pairwise comparisons performed with false-discovery-rate adjustment (31). Differences are presented with 95 % confidence intervals. In the programming of the semi-randomized assignment of the high probability reward opportunity to animals, a bug was discovered which resulted in the faulty assignment of a congruent operant corner as the high reward probability opportunity to the solo learner mice of one social group (2 animals). The respective nights (n = 9) were excluded from analysis and plotting for all animals in the study.

#### Live Mouse Tracker Data

For analysis of live mouse tracking data, scripts provided by de Chaumont *et al*. (32) were used, adapted and modified as needed. Statistical analyses on LMT data were conducted in Python (Version 3.12) and RStudio (Version 2024.04.0 Build 735). The LMT stored the tracking data in form of video recordings of 10 min length each and in SQLite databases. Further, the camera provided a three-dimensional depth scanning of the cage. For the present study, the tracking- and behavioral data in the SQLite databases was utilized. The SQLite database structure is described in detail in (22). Briefly, for each recorded frame, animal IDs, coordinates and posture are annotated as well as events (detections, RFID matches/mismatches, social behaviors). Annotated events were reconstructed and a reliability analysis of the tracking was conducted, which included calculating the rate of animals being unidentified over the total amount of recorded frames, and the rates of RFID matches and mismatches upon antenna activation.

#### Filtering of LMT Data

A method was devised to separate detection events which could be directly validated via RFID from possibly erroneous detections. Basis for operations on LMT SQLite databases were the scripts provided by de Chaumont *et al.* (32). Initially, events in the SQLite databases that could not be built during live tracking, were reconstructed according to (33). All events for which de Chaumont *et al*. provided reconstructive scripts were rebuild, with the exception of 1) Huddling, 2) WaterPoint 3) WallJump and 4) SAP. The computational definitions were adapted unaltered from (32).

The SQLite databases with reconstructed events were used for overall detection analysis on the group level. For analysis of individual animal behaviors, the following set of restrictions was applied in order to isolate the portions of the data which allowed for unambiguous assignment of animal IDs (**Fig. 3**):

1. Any movement exceeding a speed of 100 cm/s was excluded from analysis.
2. Any detection periods implying stationarity during which the center of mass of any detection mask moved less than 1 pixel between consecutive frames over a period of ≥60 s was excluded from analysis. Those restrictions were imposed on the SQLite databases directly and the ‘detection event’ label was removed from any frames which met the above criteria.
3. Of the detection events in the confines of the above-described locomotive boundaries, only those were considered which included an uninterrupted detection period including either
  a. an ‘RFID match’ event (i.e., the LMT-assigned identity was confirmed by RFID antenna activation and reading of the animal’s RFID transponder, resulting in confirmation of the previously assigned animal identity. In such cases, the entire uninterrupted detection period containing an RFID match event was labeled as ‘confirmed’ and passed down the analysis pipeline.
  b. an ‘RFID mismatch’ event (i.e., the LMT-assigned identity was revealed to be erroneous by RFID antenna activation and reading of the animal’s RFID transponder, resulting in the correction of the previously assigned animal identity. In such cases, only the uninterrupted detection period after the RFID mismatch event was labeled as ‘confirmed’ and passed down the analysis pipeline.

**Fig. 3:**
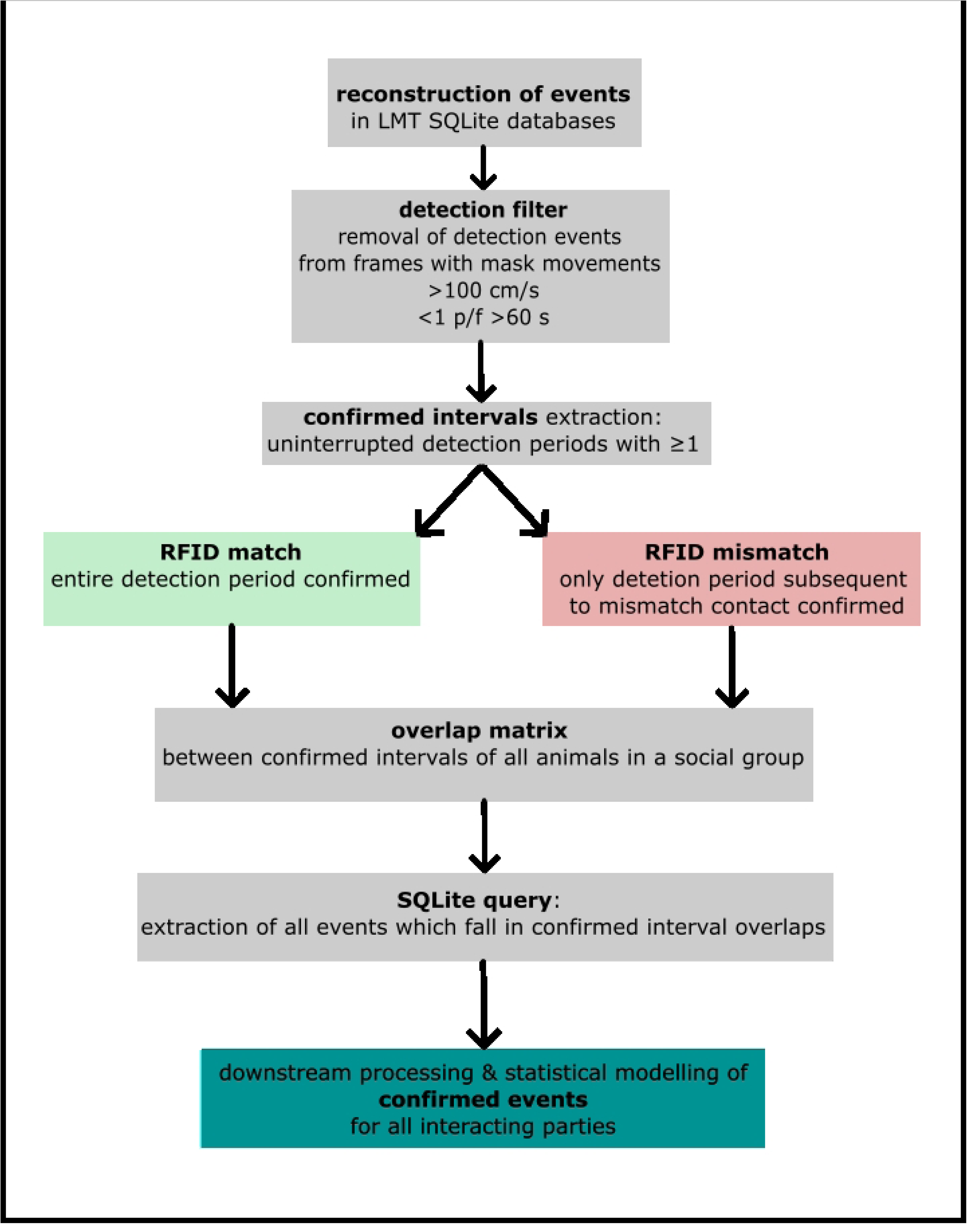
Schematic of LMT Data filtering pipeline. Events were reconstructed in the raw SQLite databases. Filtering was implemented in 2 stages: first, mask movements were limited to ≤100 cm/s and >1 pixel per consecutive frames (px/f) <60 s. Frames with mask movements of any animal exceeding these thresholds were deprived of the ‘detection’ label. In the second step, of the remaining detection periods, only those were retained which represented continuous, uninterrupted, unoccluded detections which contained at least one RFID event. For RFID match events, the entire uninterrupted detection period was confirmed, for RFID mismatch events, only the portion of the period after the events (subsequent to the ID correction) was confirmed. Of the confirmed detection periods, the overlaps across all animal combinations were identified, and any interactions of the respective combinations were extracted via SQLite query to determine the subset of interactions and behaviors of which all involved parties could be unequivocally assigned an ID.

‘Confirmed detection intervals’ were identified for each animal and subsequently, the overlapping frames of those intervals were determined for all possible combinations of animals per social group. Then, the LMT SQLite databases were queried and exclusively events which fell within these overlaps of confirmed presence between all interacting parties were considered (herein, ‘events in confirmed intervals’). In order to account for potential imbalances in confirmed detection periods, the total confirmed copresence was calculated per animal dyad and interactions were presented as interaction rates per confirmed copresence in frames.

#### Statistical modeling of dyadic interactions

For statistical modelling of dyadic social interactions, the lmerTest (30) package was deployed. Model assumptions were checked visually by Q-Q-plotting and assessing variance homogeneity of residuals versus fitted values. The model assumed average dyadic interaction density (average dyadic interaction time in frames per confirmed copresence in frames) as response variable subsequent to log-transformation to accommodate for the strictly positive right-skewed interaction density. The model assumed the fixed factors dyadic event type (20 levels) with an interaction with social learning configuration interaction type (3 levels), and social group (4 levels), as well as the interacting dyad as random intercept (24 levels). As the baseline measurement and social learning experiment differed both in length and learning tasks, we opted to model the data from both phases separately, applying the same model assumptions. This rationale was further supported by a strongly significant effect of the learning configuration × experimental phase interaction (F = 10.4; p = 3.52 × 10^-05^) when expanding the model across both phases. The overall effects of fixed factors were assessed via Type-III F-tests with Kenward–Roger degrees of freedom (30). For factors with significant effect, estimated marginal means were computed and Tukey-adjusted pairwise comparisons performed (31). Estimated means and differences are presented on the original scale with 95 % confidence intervals.

#### Correlation

For each animal, the total nightly participation (according to the conservative inclusion criteria; ref. **Fig. 3**) in LMT events ‘Approach’, ‘Approach contact’, Approach rear’, ‘Break contact’, ‘Contact’, ‘Get away’, ‘Group of 2’, ‘Move in contact, ‘Move isolated’, ‘Nest of 3’, Oral-genital contact’, Oral-oral contact‘, ‘Rear in contact’, ‘Rear isolated’, ‘Side by side contact’, ‘Side by side contact in opposite orientation’, ‘Social approach’, ‘Social escape’, ‘Stop in contact’, ‘Stop isolated’ was summed up (irrespective of interaction partners) per dark phase (19:00 – 07:00), and checked for monotonic association with the preference for the high-reward probability opportunity during the same time periods using Spearman’s rank correlation. Only behaviors were included for which ≥10 nights had both non-zero event durations and visit counts. Spearman’s Rho (herein, ϱ) was calculated for the aligned pairs via the R 4.2 base package ‘stats’(34) alongside the corresponding p-values using non-parametric percentile bootstrap (2000 resamples).

#### Code and data availability

The original python scripts for LMT analysis provided by de Chaumont *et al*. can be found at: https://github.com/fdechaumont/lmt-analysis. Any modified and original code can be found at: https://github.com/BJ-Lang/LMT-IC-Conservative-Interval-Analysis.

## 4 Results

### Preference for high-reward probability options

We demonstrated the capabilities of the LMT-IC and present the investigation of social learning in mice groups as a potential application. To that end, mice (n = 16) were presented with probabilistic reward options through the IC over a period of 41 d. During this time period, the mice were continuously tracked via the LMT component of the system. Mice were randomly assigned to either team- or solo-learning configuration, and the reward probability distribution shifted every 3 d (individually for the ‘solo learning’ mice, congruently for the ‘team learning’ mice).

**Fig. 4A** depicts the average preference for the high reward probability option of all animals in team- and solo-learning configuration over the duration of the experiment. The individual box plots represent the group average per night. Overall, co-learning (0.529 ± 0.0357) did not significantly enhance preference for the highest reward probability option over individual learning (0.501 ± 0.0358; p = 0.5982). Animals from both social learning configurations however developed a preference (28.2 ± 3.51 / 32 nights) for the high reward probability options above chance level (0.25).

**Fig. 4.**
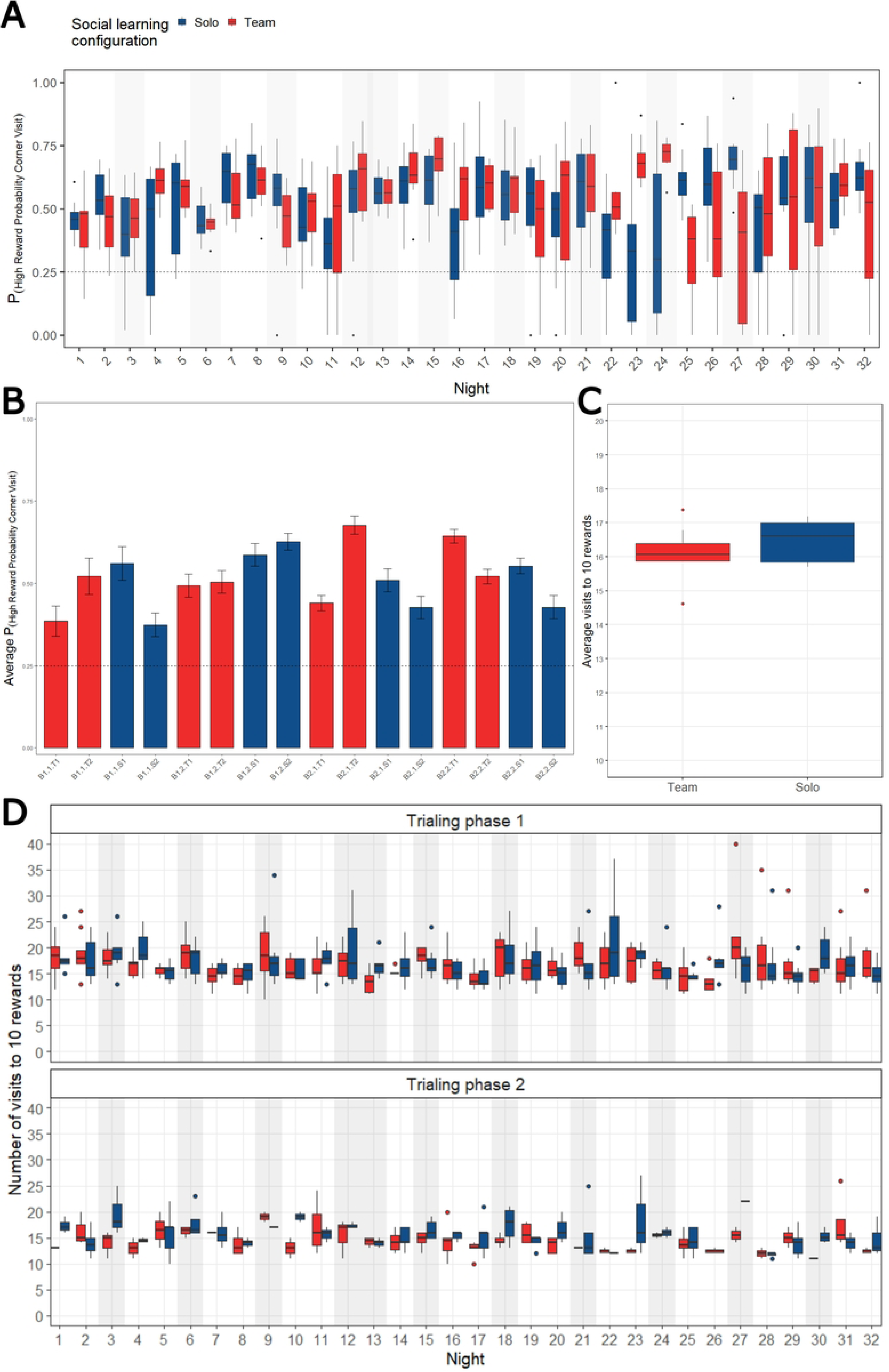
Preference for high reward probability corner (p = 0.9) in solo- and team-learning configuration. Animals continuously lived in the LMT-IC for the duration of the experiment and performed rewarded trials in 2 phases (herein, trialing phases) each night. Every third night, distribution of reward probabilities changed and the animals had to adapt. Depicted is data from 16 animals that were tested in groups of n = 4. For team learners (n = 8), the distribution of reward probabilities was congruent at any given time, whereas for solo learners, distribution of reward probabilities was different from all other animals in the group. (A) Average preference for high reward probability option over time. Y-axis depicts the average preference per night for the high-reward probability corner and the x-axis the days of the experiment. The horizontal dashed line marks chance level (i.e., one of the 4 corners). Gray underlays indicate the nights in which a change of reward probability distribution occurred. (B) Overall average preference for high reward probability corner per animal, sorted by batch (B1, B2) and social group (1, 2). Batch 1 – group 1 (B1.1), batch 1 – group 2 (B1.2), batch 2 – group 1 (B2.1) and batch 2 - group 2 (B2.2) denominate the batches / social groups. T1 and T2 denominate the team-learning animals, S1 and S2 the solo-learning animals. (C) Average number of visits per trialing phase until 10 rewards were received. (D) Number of visits until 10 rewards were received per phases 1 (21:00 – 23:00) and 2 (03:00 – 05:00) per night. Y-axis depicts the average number of visits until solo- (blue) and team-learning (red) animals collected 10 rewards *per capita*, the x-axis depicts the days of the experiment. Gray underlays indicate the nights in which a change of reward probability distribution occurred.

### Live Mouse Tracking

Over the entire duration of the experiment, animals were tracked through the LMT portion of the apparatus, which also automatically annotated individual- and social behaviors (**Fig. 5**; for a list of currently available behaviors detectable by the LMT see ref. (23)).

**Fig. 5.**
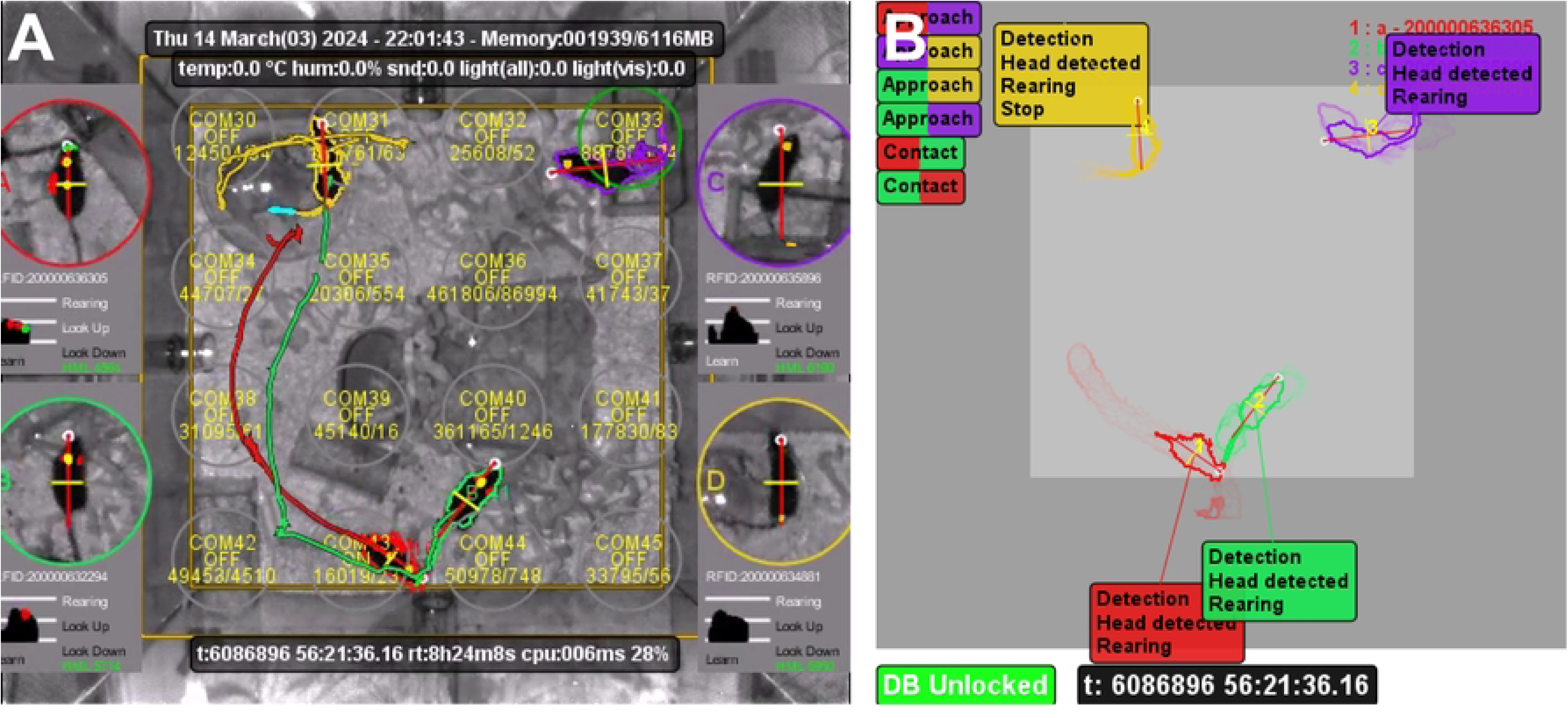
Live Mouse Tracking. (A) Top-down camera view into the cage. Capture taken 1 h after onset of the dark period, with the four mice present in the field of view. Colored masks outline the individual animals and recent trajectories. Animal 1 (red) is in contact with animal 2 (green), while animal 2 (purple) is inside the top right corner house, facing animal 4. Animal 4 (yellow) is running on the running disk. (B) Tracking view of the same timepoint as in (A). Text overlays next to animal masks detail detection and pose estimate, in the top left corner social configurations (here: approach and contact) occurring in the current frame are detailed, with the color-coding representing the animals involved.

Co-learning animals were also not significantly faster in adapting to the changing reward probabilities – however, we observed a slight non-significant trend (p = 0.1906) in team-leaning animal taking less (15.8 ± 3.76) visits to operant corners to accumulate 10 rewards in a phase than solo-learning animals (16.2 ± 4.04; **Fig. 4C-D**).

**Fig. 6** depicts the rate of animal identification during the dark phases over the entire time of the experiment, as well as the rate of RFID validations and corrections, respectively.

**Fig. 6:**
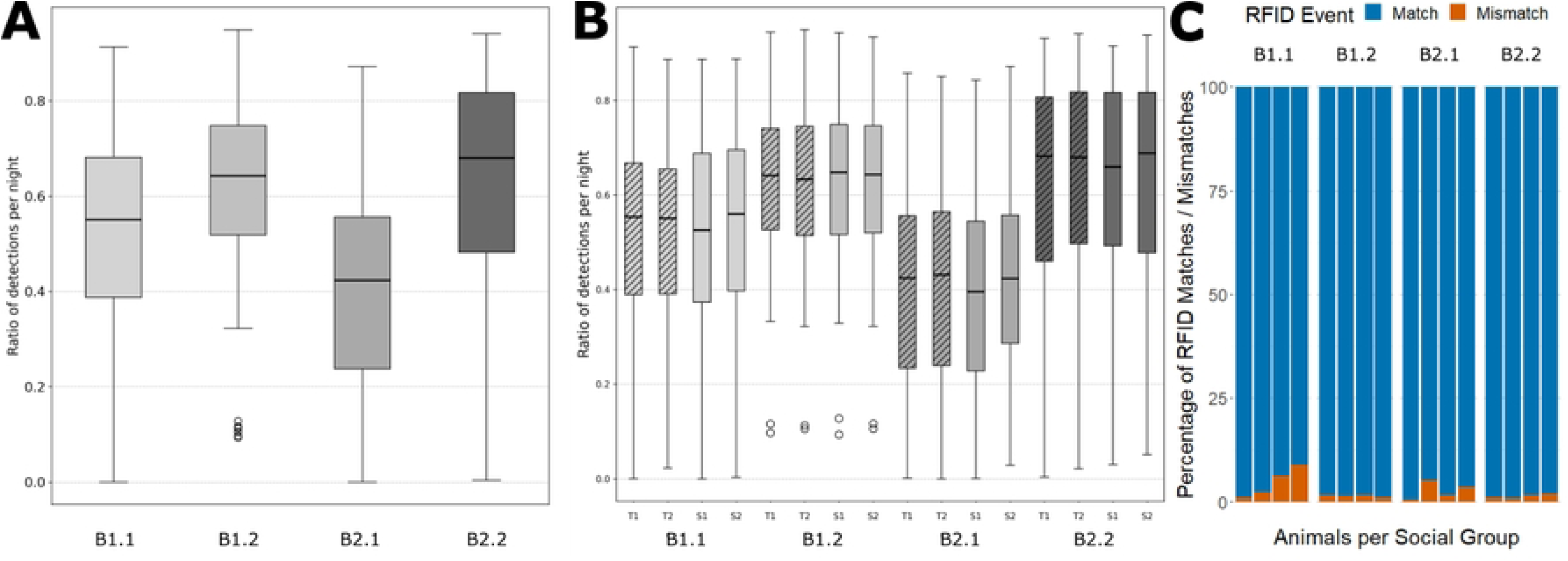
LMT detection accuracy. (A-B) Animal detection rates during dark phase (19:00-07:00). Boxplots depict the average detection rates across all nights included in the experiment per social group and animals. Each boxplot (A) and group of subplots (B), respectively, depicts data of 4 animals over the span of 75 nights (A) By social group (n = 4). (B) By individual animals (n = 16). Hatched plots represent co-learning animals, unhatched plots represent individually learning animals. (C) Percentage of RFID verification/correction (match/mismatch) during tracking. One RFID antenna at a time is always active to either identify unidentified animals, or validate the ID of presumably identified animals. The blue portion of the graph depicts the percentage of instances in which the presumed identity of either animal was validated upon antenna activation, the orange portion depicts the percentage of instances in which the animal ID was corrected upon antenna activation.

Compared to manual validation by the experimenter, automated annotation of behaviors was highly accurate. In the current configuration however, animal identification by the LMT was imperfect. Of note, the rate of dark phase animal detection here is averaged over the entire duration of the study, which also included phases of inactivity by the animals. During those, particularly after the first trialing phase ended, the animals commonly remained huddled in the nesting area, where they could not be segmented by the LMT. In those scenarios, identification could not be rescued by activation of RFID antennae, as the antenna grid resolution is not high enough to discriminate huddled stationary mice. The detection rate was highest during the first trialing phases of the night (21:00 – 23:00), when the animals were the most active (**Supporting Fig. 1**).

We attribute the imperfect animal identification mainly to the very high degree of enrichment used in the study, which lead to occlusion of animals from the camera’s field of view. The rate of RFID corrections (instances, in which an animal is supposedly identified by the tracking algorithm, but then identity-corrected upon RFID antenna activation) was low (overall 2.71 %; **Fig. 6C**) and, in fact, in a similar range of the misidentification rate reported by de Chaumont *et al*. (2.69 %; (22)) and was calculated using the same method used and provided by de Chaumont *et al*. This suggests that once animals are visible and can be segmented by the machine learning algorithm, identification is relatively stable.

Overall, mice were detected in half of all frames during the dark phases (0.543 ± 0.232), notably with the average detection rates of social groups B1.1 and B2.1 being slightly lower (0.516 ± 0.221 and 0.406 ± 0.223, respectively) than those of the two other social groups B1.2 and B2.2 (0.625 ± 0.173 and 0.626 ± 0.224, respectively). Of note, one LMT-IC system was used for groups B1.1 and B2.1, whereas another was deployed for groups B1.2 and B2.2, possibly suggesting a difference in performance in the hardware. Individual detection rates (**Fig. 7B)** mirror the detection rates on the social group level, with a considerable intergroup variance (Ø ± 0.21), but little intragroup variance (Ø ± 0.006).

**Fig. 7:**
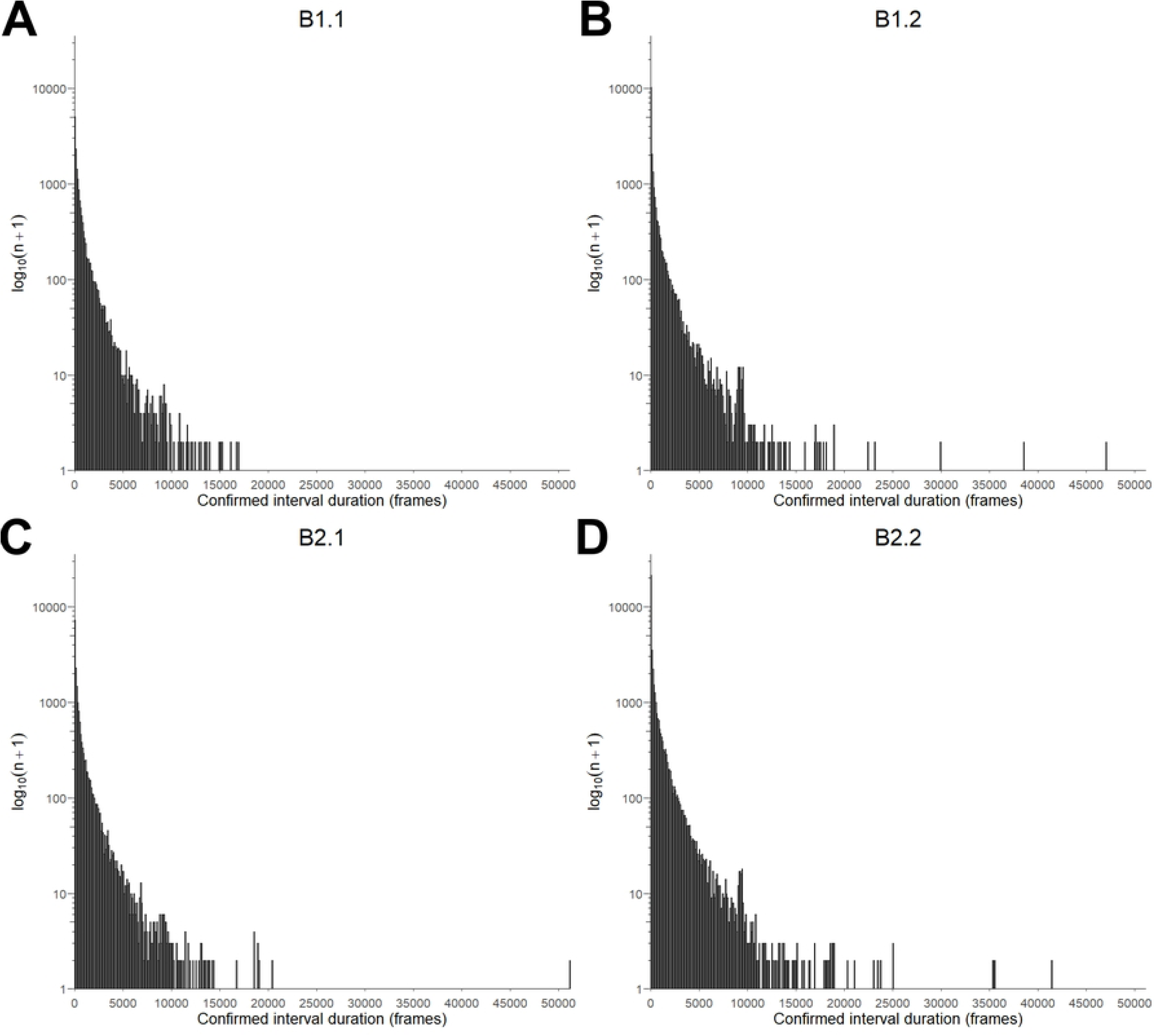
Histograms of confirmed detection intervals per group. Plots depict the frequency of detection intervals in relation to their duration, sorted by group. Frequency is expressed on a logarithmic scale for better readability. The occurrence of all intervals was increased by 1 before log transformation (log_10_(n+1)) in order to include intervals which only occurred once in the plot. (A) B1.2 (B) B1.2 (C) B2.2 (D) B2.2.

#### Misidentification in highly enriched environment

A current limitation of the LMT-IC system in a highly enriched living area is the temporary misidentification of animals which occurs when an animal is occluded by enrichment, or multiple animals frequent the same operant corner within a short time period. The high degree of enrichment deployed in our study, in concert with the opportunity for the animals to remove themselves from the camera’s field of view at any time, very likely cumbered the coherent identification of animals. This manifested as the video tracking component ‘losing’ identified animals in between RFID antenna contacts; or the assigned IDs being swapped among detection masks. De Chaumont *et al*., (22) who used a small custom-built mouse house, 5 plastic toy bricks and much less nesting material in their study as the sole enrichment, did not report such difficulties in maintaining continuous animal ID assignment.

To mitigate the misidentification of the animals, strict filters were imposed on the tracking data. For the exemplary analysis of interactions between assigned to solo- and team learning configurations, we opted for a very conservative identification method (**Fig. 3**) by only considering instances in which animal IDs could be directly confirmed by reading of the respective RFID tags - thereby omitting the majority of the data (**Table 1**).

**Table 1:** ‘Filtered’ dataset after imposing strict inclusion criteria in relation to total detection dataset.

| Group | Animal | RFID | Total detection (frames) | Confirmed detection (f) | % Confirmed |
| --- | --- | --- | --- | --- | --- |
| B1.1 | T1 | 900200000630066 | 34185438 | 1584407 | 4.63 |
|  | T2 | 900200000636022 | 33781467 | 1757181 | 5.2 |
|  | S1 | 900200000628901 | 35794953 | 3372170 | 9.42 |
|  | S2 | 900200000629604 | 37489812 | 4388775 | 11.71 |
| B1.2 | T1 | 900200000627173 | 55339945 | 3170837 | 5.73 |
|  | T2 | 900200000627861 | 56116158 | 3294259 | 5.87 |
|  | S1 | 900200000627097 | 54606962 | 2817512 | 5.16 |
|  | S2 | 900200000631863 | 54457153 | 2953909 | 5.42 |
| B2.1 | T1 | 900200000634966 | 40090609 | 2745508 | 6.85 |
|  | T2 | 900200000635982 | 38416859 | 2054144 | 5.35 |
|  | S1 | 900200000633186 | 39157686 | 2043219 | 5.22 |
|  | S2 | 900200000635944 | 42647717 | 4539567 | 10.64 |
| B2.2 | T1 | 900200000634881 | 59358641 | 3832393 | 6.46 |
|  | T2 | 900200000636305 | 60543233 | 5373509 | 8.88 |
|  | S1 | 900200000632294 | 61287844 | 6775174 | 11.05 |
|  | S2 | 900200000635896 | 59918622 | 5650570 | 9.43 |

The ‘confirmed detection intervals’ ranged from 1 - 51173 frames in length, with the distribution being heavily skewed towards short intervals (**Fig. 7**). This was, to a degree, accounted for when selecting the linear mixed model assuming the interaction density per confirmed copresence as response variable. However, the filtering process inherently favored shorter interactions, with the probability that a ‘confirmed’ detection period remaining uninterrupted by identification errors decaying exponentially with interval length. For all subsequent analyses, the total interaction durations of all occurrences of an interaction type per time unit were considered rather than the durations of individual interactions.

#### Dyadic interaction types

To investigate whether co- or individually learning animals differed in their social interactions, we fit a linear mixed-effects model to the log-transformed interaction rates per dyad. We observed a highly significant effect of the interaction type on the response variable (**Fig. 8A**; F = 124.149; p < 2.2 × 10^-16^). The by far most commonly observed dyadic interaction was animals approaching each other, which occurred almost every other frame of confirmed copresence (0.499 ± 0.0726). This was however true for all animals regardless of learning configuration (T-T: 0.497 ± 0.1449; T-S: 0.499 ± 0.0727; S-S: 0.499 ± 0.1454). Particularly events which were defined by prolonged, more complex interactions were much less common, such as for example long chase (0.003 ± 3.72 × 10^-4^), and sequences of oral-oral contact alternating to oral-genital contact (0.011 ± 1.28 ×10^-3^), as well as the inverse (0.009 ± 1.01 × 10^-3^), respectively. This may of course in part be facilitated by the LMT detection interval filtering rationale favoring shorter detection intervals, and conversely (**Fig. 7**), shorter dyadic interactions. However, these complex interactions were also observed more often in the pre-processed annotated tracking data, suggesting an inherently lower rate.

**Fig. 8:**
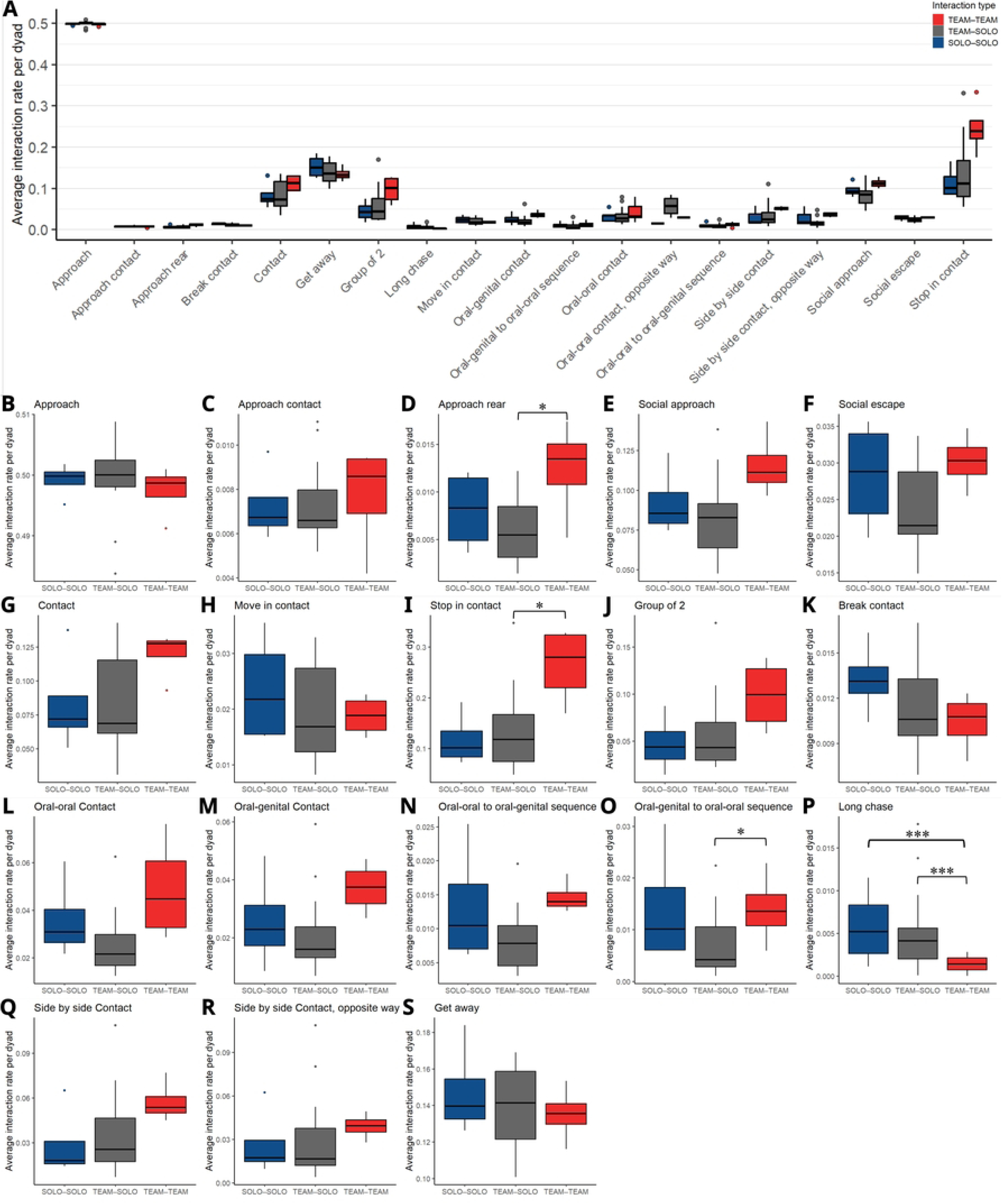
Average dyadic interaction rates per confirmed copresence frame. Boxplots depict average dyadic interaction type. To minimize ID inaccuracies, only tracks were considered that 1) contained at least 1 successful RFID read and 2) were within the maximum speed limit (100 cm/s) as well as 3) above the movement threshold (1 pixel per consecutive frames, herein px/f, for >60 s). These criteria had to be met by all interaction partners in order for the event to be considered. Lines represent median; lower hinges represent 0.25 quantiles; upper hinges represent 0.75 quantiles; whiskers represent ≤ Q3 + 1.5 × IQR and ≥ Q1 − 1.5 × IQR, respectively. Color-coding represents the categories of interaction between social learning configurations: solo-solo (blue), team - solo (gray) and team-team (red). Y-axes represent the average interaction rates per dyad: the total interaction durations in frames for all event types were calculated per dyad; as well as the confirmed copresence in frames for all dyads. The ratio of total interaction duration / confirmed copresence per dyad represents the interaction rate. Asterisks represent confidence levels: *: p ≤ 0.05, **: p ≤ 0.01, ***: p ≤ 0.001. (A) Overview plot. Along the x-axis, the observed dyadic interactions were plotted on a uniform linear y-axis. (B-S) Individual plots per interaction type with individual y-scales for comparison of the social learning configuration classes: (B) Approach (C) Approach contact (D) Approach rear (E) Social approach (F) Social escape (G) Contact (H) Move in contact (I) Stop in contact (J) Group of 2 (K) Break contact (L) Oral-oral contact (M) Oral-genital contact (N) Oral-oral to oral-genital contact sequence (O) Oral-genital to oral-oral contact sequence (P) Long chase (Q) Side-by-side contact (R) Side-by-side contact in opposite orientation (S) Get away.

Social learning configuration in itself did not have a significant effect on the average dyadic interaction duration (F = 2.755, p = 0.0887), however, crucially, the social learning configuration had a significant effect on the rates of certain interaction types, as evidenced by the significant interaction of event type × learning configuration (**Fig. 8B-S**; F = 2.04, p = 0.0005).

Co-learning animals approached more often rearing teammates than rearing solo-learners (T-T: 0.011 ± 0.0033 vs. T-S: 0.005 ± 0.0007; p = 0.0316) and stopped more often while being in contact with a teammate than with an individually learning animal (T-T: 0.255 ± 0.074 vs. T-S: 0.116 ± 0.0169; p = 0.043). Further, co-learning animals transitioned more frequently from oral-genital contact to oral-oral contact with their teammates than with solo-learners (T-T: 0.013 ± 0.0037 vs. T-S: 0.005 ± 0.0008; p = 0.0233). Intriguingly, all the interactions co-learning animals engaged more often in than their cagemates are associated with pro-social behavior, whereas the individually learning mice participated more often in ‘long chases’ among each other (S-S: 0.0041 ± 0.0012 vs. T-T: 0.0003 ± 0.0001; p < 0.0001) and with team-learning animals than those did among each other (S-T: 0.0028 ± 0.0004 vs. T-T: 0.0003 ± 0.0001; p < 0.0001). Chasing behavior in female rodents is usually associated with aggressive or antagonistic behavior (35). During the initial baseline measurement of learning performance, there were no significant effects of the future randomly assigned social learning configuration (F = 1.83; p = 0.1880) or the interaction of prospective social learning configuration × interaction type on the dyadic interaction rates (F = 1.05; p = 0.3989; **Supporting Fig. 2**).

Of note, these observations were made on a heavily restricted subset of all recording data. While the vast amount of tracking data still allowed us to observe differences in some behaviors between individually- and co-learning animals, we therefore refrain from drawing final conclusions on the nature of social learning in mice within the scope of this study. Nevertheless, this proof-of-concept study demonstrates the ability of the method presented here to detect subtle nuances in social interactions in a fully automated manner even in subpar environments and noisy datasets.

#### Correlation of individual preference for high-reward option with social behavior

With animals belonging to either learning configuration differing in some behaviors, we proceeded to probe whether learning performance (i.e., preference for the high reward probability option) was linked to individual- or social behaviors on the individual level. To this end, we checked for monotonic association between the nightly individual learning performance and different behaviors which could be linked to prosocial- or socially aversive interactions (**Table 2**) using Spearman’s ranked correlation. Significant correlation between preference for the high reward probability option and social interactions was only observed sporadically, with the strongest correlation being established between the preference for the highest reward probability option for one individually learning animal (B1.1.S2) and ‘rearing in contact’ (ϱ = 0.52, p = 0.0033), which arguably represent a socio-positive interaction. Another strong correlation was observed for a team-learning animal of the same social group (B1.1.T1) between preference for the high reward probability corner and contact with other animals (ϱ = 0.5, p = 0.0045). For this animal, we observed the most associations between learning performance and socio-positive behaviors (approach: ϱ = 0.36, p = 0.0443; approach rear: ϱ = 0.44, p = 0.0129; contact: ϱ = 0.5, p = 0.0045; move in contact: ϱ = 0.44, p = 0.014; oral-oral contact: ϱ = 0.38, p = 0.0353; side-by-side contact: ϱ = 0.42, p = 0.0198; side-by-side contact in opposite orientation: ϱ = 0.45, p = 0.0113; social approach: ϱ = 0.41, p = 0.0213; and stop in contact: ϱ = 0.41, p = 0.0231 – but also with aversive behaviors (rear isolated: ϱ = 0.39, p = 0.0296 and social escape: ϱ = 0.39, p = 0.0291). Overall, we found a monotonic association between nightly learning performance and 11/20 probed behaviors for animal B1.1.T1, hinting towards a possible relationship between social interactions and learning success for this animal in the co-learning paradigm.

**Table 2:** Assessment of monotonic association between individual learning performance and different social behaviors using Spearman’s ranked correlation. The total durations per night of automatically annotated social interactions were correlated with the nightly preference for the high reward probability option for all individual animals in four social groups (B1.1, B1.2, B2.1, B2.2). Animals assigned to co-learning configuration are labeled T1 and T2; individual learning configuration are labeled S1 and S1. n represents the number of learning- and behavioral data pairings, which was derived from the number of nights during which a behavior could be confirmed by according to Fig. 3. ϱ represents Spearman’s Rho with the p-value p (95 % bootstrap-CI). Statistically significant correlations are depicted in bold.

|  |  | B1.1 |  |  |  | B1.2 |  |  |  | B2.1 |  |  |  | B2.2 |  |  |  |
| --- | --- | --- | --- | --- | --- | --- | --- | --- | --- | --- | --- | --- | --- | --- | --- | --- | --- |
|  |  | T1 | T2 | S1 | S2 | T1 | T2 | S1 | S2 | T1 | T2 | S1 | S2 | T1 | T2 | S1 | S2 |
| Approach | n | 31 | 31 | 31 | 30 | 32 | 32 | 32 | 32 | 32 | 32 | 32 | 32 | 32 | 32 | 32 | 32 |
| | $\rho$ | **0.36** | 0.01 | -0.17 | **0.37** | -0.06 | -0.16 | -0.11 | -0.05 | -0.20 | -0.28 | 0.11 | 0.32 | 0.12 | 0.17 | -0.03 | 0.24 |
|  | p | **0.04** | 0.98 | 0.36 | **0.05** | 0.76 | 0.38 | 0.56 | 0.81 | 0.27 | 0.13 | 0.55 | 0.08 | 0.53 | 0.37 | 0.88 | 0.19 |
| Approach contact | n | 31 | 31 | 31 | 30 | 32 | 32 | 32 | 32 | 32 | 32 | 32 | **32** | 32 | 32 | 32 | 32 |
| | $\rho$ | 0.32 | 0.06 | -0.16 | 0.30 | -0.14 | -0.26 | -0.23 | 0.08 | -0.15 | -0.15 | 0.12 | **0.35** | -0.04 | 0.11 | -0.05 | -0.11 |
|  | p | 0.08 | 0.75 | 0.39 | 0.10 | 0.46 | 0.14 | 0.21 | 0.68 | 0.42 | 0.40 | 0.52 | **0.05** | 0.83 | 0.54 | 0.79 | 0.57 |
| Approach rear | n | 31 | 31 | 31 | 30 | 32 | 32 | 32 | 32 | 32 | 32 | 32 | 32 | 32 | 32 | 32 | 32 |
| | $\rho$ | **0.44** | 0.06 | -0.15 | **0.40** | -0.09 | -0.15 | -0.27 | 0.17 | -0.13 | -0.02 | 0.14 | 0.08 | -0.02 | 0.08 | -0.13 | -0.14 |
|  | p | **0.01** | 0.75 | 0.42 | **0.03** | 0.64 | 0.43 | 0.14 | 0.37 | 0.47 | 0.90 | 0.45 | 0.68 | 0.92 | 0.68 | 0.49 | 0.44 |
| Break contact | n | 31 | 31 | 31 | 30 | 32 | **32** | 32 | 32 | 32 | 32 | 32 | **32** | 32 | 32 | 32 | 32 |
| | $\rho$ | 0.26 | 0.02 | -0.10 | **0.44** | -0.16 | **-0.37** | -0.18 | -0.15 | -0.22 | -0.15 | 0.16 | **0.36** | 0.04 | 0.14 | -0.05 | 0.13 |
|  | p | 0.16 | 0.91 | 0.61 | **0.02** | 0.37 | **0.03** | 0.32 | 0.40 | 0.23 | 0.42 | 0.37 | **0.04** | 0.83 | 0.46 | 0.78 | 0.49 |
| Contact | n | 31 | 31 | 31 | 30 | 32 | 32 | 32 | 32 | 32 | 32 | 32 | 32 | 32 | 32 | 32 | 32 |
| | $\rho$ | **0.50** | -0.12 | -0.14 | 0.30 | 0.07 | -0.21 | -0.21 | -0.13 | -0.16 | -0.22 | 0.15 | 0.10 | 0.03 | 0.02 | -0.01 | 0.06 |
|  | p | **.005** | 0.51 | 0.44 | 0.11 | 0.71 | 0.24 | 0.26 | 0.48 | 0.39 | 0.24 | 0.41 | 0.61 | 0.89 | 0.92 | 0.94 | 0.75 |
| Get away | n | 31 | 31 | 31 | 30 | 32 | 32 | 32 | 32 | 32 | 32 | 32 | 32 | 32 | 32 | 32 | 32 |
| | $\rho$ | 0.31 | 0.00 | -0.13 | 0.35 | -0.15 | -0.24 | -0.21 | 0.24 | -0.18 | -0.24 | 0.00 | 0.26 | -0.06 | 0.14 | -0.12 | -0.20 |
|  | p | 0.09 | 0.99 | 0.49 | 0.06 | 0.42 | 0.19 | 0.24 | 0.18 | 0.33 | 0.19 | 0.99 | 0.15 | 0.74 | 0.44 | 0.50 | 0.29 |
| Group of 2 | n | 31 | 31 | 31 | 30 | 32 | 32 | **32** | 32 | 32 | 32 | 32 | 32 | 32 | 32 | 32 | 32 |
| | $\rho$ | 0.34 | -0.19 | -0.11 | 0.32 | 0.06 | -0.33 | **-0.35** | 0.06 | -0.22 | -0.13 | 0.10 | 0.20 | -0.04 | 0.20 | -0.12 | -0.21 |
|  | p | 0.06 | 0.31 | 0.57 | 0.09 | 0.76 | 0.06 | **0.05** | 0.76 | 0.23 | 0.50 | 0.58 | 0.28 | 0.82 | 0.28 | 0.52 | 0.25 |
| Move in contact | n | 31 | 31 | 31 | 30 | 32 | **32** | 32 | 32 | 32 | 32 | 32 | 32 | 32 | 32 | 32 | 32 |
| | $\rho$ | **0.44** | 0.05 | -0.02 | **0.41** | -0.07 | **-0.44** | -0.29 | 0.07 | -0.21 | -0.10 | 0.13 | 0.14 | 0.20 | 0.04 | -0.04 | 0.06 |
|  | p | **0.01** | 0.81 | 0.92 | **0.02** | 0.72 | **0.01** | 0.11 | 0.70 | 0.24 | 0.59 | 0.49 | 0.44 | 0.28 | 0.84 | 0.85 | 0.73 |
| Move isolated | n | 31 | 31 | 31 | 30 | 32 | 32 | 32 | 32 | 32 | 32 | 32 | 32 | 32 | 32 | 32 | 32 |
| | $\rho$ | 0.23 | -0.12 | -0.05 | 0.33 | -0.12 | -0.19 | -0.21 | 0.30 | -0.12 | -0.22 | -0.06 | 0.12 | 0.20 | 0.20 | -0.05 | 0.19 |
|  | p | 0.21 | 0.54 | 0.79 | 0.08 | 0.51 | 0.30 | 0.26 | 0.10 | 0.53 | 0.23 | 0.73 | 0.53 | 0.27 | 0.28 | 0.80 | 0.30 |
| Nest of 3 | n | 31 | 31 | 31 | 30 | 32 | 32 | 32 | 32 | 32 | 32 | 32 | 32 | 32 | **32** | 32 | 32 |
| | $\rho$ | 0.10 | -0.02 | -0.26 | 0.25 | 0.03 | -0.19 | -0.07 | 0.21 | -0.22 | 0.08 | -0.05 | 0.01 | 0.15 | **0.36** | -0.05 | -0.27 |
|  | p | 0.58 | 0.90 | 0.16 | 0.18 | 0.86 | 0.30 | 0.70 | 0.26 | 0.24 | 0.68 | 0.81 | 0.94 | 0.43 | **0.04** | 0.78 | 0.13 |
| Oral-genital contact | n | 31 | 31 | 31 | 30 | 32 | 32 | 32 | 32 | 32 | 32 | 32 | 32 | 32 | 32 | 32 | 32 |
| | $\rho$ | 0.33 | -0.13 | -0.16 | 0.26 | -0.11 | -0.34 | -0.24 | 0.09 | -0.24 | 0.10 | 0.04 | 0.15 | 0.06 | -0.12 | 0.03 | -0.02 |
|  | p | 0.07 | 0.49 | 0.41 | 0.17 | 0.53 | 0.06 | 0.18 | 0.62 | 0.19 | 0.59 | 0.84 | 0.41 | 0.75 | 0.51 | 0.87 | 0.94 |
| Oral-oral contact | n | 31 | 31 | 31 | 30 | 32 | 32 | 32 | 32 | 32 | 32 | 32 | 32 | 32 | 32 | 32 | 32 |
| | $\rho$ | **0.38** | 0.15 | -0.11 | 0.36 | 0.14 | -0.21 | -0.14 | 0.07 | -0.13 | -0.31 | 0.16 | 0.18 | 0.10 | 0.02 | 0.02 | -0.02 |
|  | p | **0.04** | 0.41 | 0.56 | .053 | 0.44 | 0.24 | 0.46 | 0.72 | 0.46 | 0.09 | 0.39 | 0.33 | 0.60 | 0.90 | 0.90 | 0.94 |
| Rear in contact | n | 31 | 31 | 31 | 30 | 32 | **32** | 32 | 32 | 32 | 32 | 32 | 32 | 32 | 32 | 32 | 32 |
| | $\rho$ | 0.32 | 0.07 | -0.11 | **0.52** | -0.21 | **-0.42** | -0.16 | 0.18 | -0.06 | -0.29 | 0.22 | 0.21 | -0.08 | 0.19 | -0.07 | -0.11 |
|  | p | 0.08 | 0.71 | 0.56 | **.003** | 0.24 | **0.02** | 0.40 | 0.33 | 0.75 | 0.11 | 0.24 | 0.25 | 0.66 | 0.30 | 0.71 | 0.55 |
| Rear isolated | n | 31 | 31 | 31 | 30 | 32 | 32 | 32 | 32 | 32 | 32 | 32 | 32 | 32 | 32 | 32 | 32 |
| | $\rho$ | **0.39** | 0.01 | -0.14 | **0.38** | -0.12 | -0.07 | -0.14 | 0.17 | -0.23 | -0.28 | 0.09 | 0.08 | 0.05 | 0.34 | 0.07 | -0.32 |
|  | p | **0.03** | 0.94 | 0.46 | **0.04** | 0.52 | 0.69 | 0.43 | 0.35 | 0.21 | 0.13 | 0.63 | 0.66 | 0.80 | 0.06 | 0.70 | 0.07 |
| Side by side contact | n | 31 | 31 | 31 | 30 | 32 | 32 | 32 | 32 | 32 | 32 | 32 | 32 | 32 | 32 | 32 | 32 |
| | $\rho$ | **0.42** | -0.14 | -0.14 | 0.33 | 0.22 | -0.18 | -0.18 | -0.03 | -0.11 | -0.13 | -0.14 | 0.06 | -0.02 | -0.19 | -0.06 | -0.03 |
|  | p | **0.02** | 0.47 | 0.46 | 0.08 | 0.23 | 0.32 | 0.32 | 0.87 | 0.56 | 0.47 | 0.44 | 0.75 | 0.90 | 0.29 | 0.75 | 0.86 |
| Side by side, opposite | n | 31 | 31 | 31 | 30 | 32 | 32 | 32 | 32 | 32 | 32 | 32 | 32 | 32 | 32 | 32 | 32 |
| | $\rho$ | **0.45** | 0.06 | -0.19 | 0.28 | 0.15 | -0.16 | -0.24 | -0.07 | -0.10 | -0.13 | 0.32 | 0.19 | -0.09 | -0.08 | 0.00 | 0.03 |
|  | p | **0.01** | 0.77 | 0.31 | 0.13 | 0.42 | 0.40 | 0.19 | 0.71 | 0.59 | 0.46 | 0.07 | 0.30 | 0.65 | 0.65 | 1.00 | 0.89 |
| Social approach | n | 31 | 31 | 31 | 30 | 32 | 32 | 32 | 32 | 32 | 32 | 32 | 32 | 32 | 32 | 32 | 32 |
| | $\rho$ | **0.41** | -0.15 | -0.12 | 0.34 | -0.06 | -0.27 | -0.17 | 0.12 | -0.26 | -0.12 | 0.06 | 0.18 | -0.20 | 0.06 | -0.14 | -0.19 |
|  | p | **0.02** | 0.41 | 0.51 | 0.07 | 0.76 | 0.13 | 0.36 | 0.51 | 0.15 | 0.51 | 0.75 | 0.32 | 0.27 | 0.73 | 0.43 | 0.30 |
| Social escape | n | 31 | 31 | 31 | 30 | 32 | **32** | 32 | 32 | 32 | 32 | 32 | 32 | 32 | 32 | 32 | 32 |
| | $\rho$ | **0.39** | -0.09 | -0.04 | **0.38** | -0.15 | **-0.36** | -0.21 | 0.19 | -0.18 | -0.03 | 0.16 | 0.16 | -0.07 | 0.06 | -0.14 | -0.17 |
|  | p | **0.03** | 0.64 | 0.81 | **0.04** | 0.41 | **0.04** | 0.26 | 0.29 | 0.32 | 0.86 | 0.38 | 0.39 | 0.71 | 0.73 | 0.44 | 0.36 |
| Stop in contact | n | 31 | 31 | 31 | 30 | 32 | 32 | 32 | 32 | 32 | 32 | 32 | 32 | 32 | 32 | 32 | 32 |
| | $\rho$ | **0.41** | -0.10 | -0.17 | 0.28 | -0.02 | -0.19 | -0.26 | 0.03 | -0.19 | -0.19 | 0.14 | 0.15 | -0.17 | 0.09 | -0.08 | -0.22 |
|  | p | **0.02** | 0.59 | 0.37 | 0.14 | 0.90 | 0.30 | 0.14 | 0.88 | 0.29 | 0.30 | 0.44 | 0.41 | 0.36 | 0.62 | 0.65 | 0.22 |
| Stop isolated | n | 31 | 31 | 31 | 30 | 32 | 32 | 32 | 32 | 32 | 32 | 32 | 32 | 32 | 32 | 32 | 32 |
| | $\rho$ | 0.19 | 0.00 | -0.15 | **0.40** | -0.12 | -0.25 | -0.08 | 0.14 | -0.12 | -0.17 | 0.13 | -0.01 | -0.04 | 0.20 | -0.14 | -0.21 |
|  | p | 0.30 | 1.00 | 0.42 | **0.03** | 0.51 | 0.17 | 0.65 | 0.46 | 0.52 | 0.35 | 0.49 | 0.97 | 0.82 | 0.26 | 0.44 | 0.25 |

However, this observation did not hold true for the other team-learning animals. For the remaining mice assigned to co- learning, only sporadic correlation between nightly preference for the best reward option and social behaviors was observed. For animal B1.2.T2, four behaviors were associated with nightly learning performance – albeit, strikingly, as opposed as for animal B1.1.T1, all of them negatively (break contact: ϱ = - 0.37, p = 0.0348; move in contact: ϱ = -0.44, p = 0.0128; rear in contact ϱ = -0.42, p = 0.0162; social escape: ϱ = -0.357, p = 0.0448). We did not find any association between social interaction and learning performance for any of the other team-learning animals, except for one behavior in B2.2.T2 (nest of 3: ϱ = 0.36, p = 0.0438). Beyond that, sporadic monotonic associations between learning and social interactions were observed for the animals assigned to individual learning, with only B1.1.S2 standing out with 8/20 probed behaviors correlated to nightly learning performance (approach: ϱ = 0.37, p = 0.0474; approach rear: ϱ = 0.4, p = 0.0284; break contact: ϱ = 0.44, p = 0.0154; move in contact: ϱ = 0.41, p = 0.0235; rear in contact: ϱ = 0.52, p = 0.0033; as well as rear in isolation: ϱ = 0.38, p = 0.0393). Crucially however, the animal with by far the most associations between socio-positive behaviors and learning performance (team-learning animal B1.1.T1) expressed, in fact, the penultimate overall preference for the best reward option (**Fig. 4B**; 0.385 ± 0.254). Only the individually learning animal B1.1.S2 – for which a monotonic association was found for 8/20 behaviors - demonstrated a lower learning performance (0.374 ± 0.196). The observation of numerous monotonic associations between nightly social interactions and preference for the best reward option only in the two ‘worst-performing’ animals may suggest socio-positive behavior to actually counteract learning success. Of note however, as we did not observe a negative correlation with such behaviors in the animals with the highest learning performances, this more likely reflects individuality in the animals rather than a general effect; and should be interpreted with caution due to the restrictive criteria of behavioral data inclusion.

Overall, no consistent association between learning performance and any individual behavior or social learning configuration could be established, suggesting learning success in this social learning paradigm to not be solely dependent on individual interaction types.

In conclusion, we here demonstrated a nearly fully automated method for presenting place learning tasks to socially-housed mice in a superenriched environment, while recording and annotating all their behaviors. Behavioral annotation was highly reliable, whereas the individual identification was severely impacted for the substantial degree of visual occlusion from the video tracking component. We mitigated this by only considering an exemplary subset of the tracking data for the present analysis, which was achieved by applying several layers of filters on visual tracking data, so that only those detection intervals and automatically annotated behaviors within those detection intervals remained for which a direct link to an ID confirmation via RFID could be established. This method proved highly reliable, but allowed us to only consider a small subsection of the behavioral data. Due to the vast amount of data gathered by the continuous animal tracking over the course of weeks, this still allowed us to make profound observations on animal interactions – yet, we strongly suggest future studies with a similar setup to rethink the amount and type of enrichment and opportunity for occlusion to ensure more reliable animal identification.

We exemplarily applied this method to a probabilistic social learning paradigm, during which reward probabilities continuously shifted and had to be relearned by the animals, either individually or in ‘teams’. We found the co-learning setting to not facilitate learning performance for the animals, but demonstrated co-learning animals to engage more often in some prosocial interactions – i.e., approaching rearing animals, stopping while in contact, and oral-genital to oral-oral contact sequences – than individually learning animals. On the other hand, individually learning animals chased other solo-learners more often than team-learners, or than co-learning animals chased each other. Overall, interactions could only sporadically be correlated to learning performance. Rather, we here established a framework which, after some careful refinement, may shed further light on the nature of learning in social interactions in mice in future studies.

## 5 Discussion

Here, we present a method which utilized a widely-used commercial system for automated behavioral testing (IntelliCage (20)) and a cost-effective, easy-to-built open-source solution for 24/7 animal tracking as well as behavioral annotation (Live Mouse Tracker (22)) to achieve comprehensive investigation of social learning and behaviors in mice. Combining the two apparatuses into one system is achievable by straightforward modifications, with the main concern being reducing RFID interference to a level allowing RFID antennae in both apparatuses to achieve reliable animal identification. Here, we did this by simply placing the IC operant corners outside the living area and connecting them to the cage via PMMA tubes of 40 cm length. Animals were observed to enter the tubes readily and willingly, with no apparent increased aversion to participate in trials in the operant corners compared to the standard IC configuration with the operant corners placed inside the cage area.

The present paper represents a report on a method which still has opportunity for further optimization. While we could observe some differences in social interactions between animals which were assigned to co-learning and individual learning paradigms, those did not manifest in significant differences in learning success. We anticipate the full potential of the method to only be realized once outstanding obstacles in animal identification reliability are overcome.

The greatest current limitation of the system lies in the identification accuracy of the live tracking component in a heavily enriched area as employed in the present study which allows the animals to disappear from the camera’s field of view. For once, this manifests as subpar detection rates (rate = 0.543 ± 0.232 overall during dark phases). Of note, we observed slightly higher overall detection rates for two of the social groups (B1.2 and B2.2) than for the other social groups (B1.1 and B2.1). The grouping happens to correlate with the LMT-IC systems deployed in the study. While this may be coincidental, and it is conceivable the social groups differed in hiding behavior and therefore occlusion from the camera, this observation suggests one of the hardware systems to perform more reliably in tracking than the other. At this time, we cannot offer a conclusive explanation for this observation, as both systems were constructed to the same standard. Irrespective of the ultimate cause, this highlights the need of precise antenna tuning and prolonged testing of the system accuracy and functionality even during ongoing studies.

Most severely impacted by the animal occlusion however was the animal identification rate. Without all animals being present in the LMT’s field of view at the same time, animal ID proved to be highly unreliable, with tracking masks commonly flipping between animals. Only when an antenna contact was made long enough for the RFID tag to be read, IDs could be reliably restored. Notably, de Chaumont *et al*. 2019, who reported a much higher rate of correct animal ID assignment (97.31 %), in their study deployed no enrichment beyond 1) wooden bedding, 2) 5 plastic toy bricks, 3) 6 cotton rolls, 4) a few brown paper strips, and 5) a monobloc mouse house consisting of one bend piece of PMMA (22). The latter was open on two sides and, by virtue of being of red PMMA, which is entirely transmissible to IR light, did not obstruct animal tracking in any way. Furthermore, no material sufficient to build a full nest (as, for instance, paper towels) was provided and in contrast to the present study, the animals had no opportunity to leave the arena at any time. Hence, all four animals per group were visible to the camera at all times, with no opportunity for any occlusion.

This setup is suitable to house groups of mice for as short time periods as carried out by de Chaumont *et al*. (max. 3 d) (22). In contrast, in the present study, which was carried out at the facilities of the German Center for the Protection of Laboratory Animals (Bf3R (36)), the animals were housed in the LMT-IC apparatus for a total of 75 d (including all habituation, baseline measurement, and experimental phases combined). This demanded a much higher standard of housing including enrichment to ensure animal welfare over extended time periods. As a result, we applied a preexisting live tracking algorithm to a much more dynamic and enriched environment than had been tested before (22,37). This superenriched environment, in which each animal also disappeared on average ∼50× /d from the trackable area, presented a far greater challenge for the LMT component than was present in the de Chaumont *et al*. study, highlighting the current limits of the method.

Being confronted with the animal identification issues, we initially strived to reconstruct misidentified animal tracks. As in contrast to the video identification, RFID readings proved highly reliable, we used any RFID antenna readings as fixed points of confirmed animal identity in space and time. From these ‘anchor points’, which we could confidently link to an animal’s identity, we computationally moved through the tracking data bidirectionally frame by frame and checked whether the animal trajectories assumed by the LMT algorithm could in fact represent natural movements (i.e., animals not exceeding a distance of 50 p ≙ 8.62 cm between consecutive frames at 30 fps) of the animals. The LMT itself has no such safeguarding mechanism and will assign animal IDs solely based on the probability based on its computer vision component, and attempt to validate them by RFID antenna activation (22). In the presence of occlusion, this can lead to animal tracking masks flipping between animals, and the apparent ‘teleportation’ of animals across the arena. We strived to computationally reconstruct actual animal trajectories in such cases by completely disregarding the LMT-assigned IDs between RFID antenna contacts, and re-assigning the IDs based on a probability matrix calculating the most likely trajectories from the confirmed antenna readings while restricting any ‘impossible’ animal movements.

The reconstructive efforts proved successful in instances when animals were at some distance from each other. In instances when animals were interacting closely, and/or under occlusion, the reconstructive restoration of ID assignments failed. Hence, we opted here to rely only on the ‘filtered’ dataset, which fulfilled strict criteria of ID certainty (ref. **Fig. 3**). The reconstructive approach will require considerably more development before being able to be deployed reliably – and the sensibility of dedicating further time and resources to bring the reconstructive approach to fruition stands to question, as in future studies, reliability in LMT tracking to a certain degree could be achieved by considerably more straightforward and easy methods, namely: 1) decreasing the degree of enrichment to prevent occlusion of the animals by enrichment components (22) or 2) using specialized enrichment and nesting material which appears opaque to the murine vision (38) but is penetrable by infrared light.

Reliable automated live multi-animal tracking continues to present a challenge for researchers – especially, when no external markings on the animals are deployed. Pose estimators such as DeepLabCut (11) achieved considerable reliability and widespread use (39), but require manual training. Very recently, a promising new approach was presented with the so-called ‘Tailtag’, a simple 3D-printable ArUco code which can be attached to mouse tails and is reported to allow for reliable tracking in social groups of up to 20 animals over the period of 7 d, outperforming the LMT in its multiplexing capabilities (40). However, this Tailtag still cannot overcome the problem presented by visual occlusion from the camera view by enrichment or structural components of the experimental setup such has trialing corners, and unlike the LMT, relies on external markings of the animals – albeit, less invasive ones as for instance bleaching of the fur (37). We anticipate future developments in the field of automated multi-animal with excitement and hope to offer a modest perspective on the current capabilities and limitations of markerless methods in highly enriched environments over prolonged time periods.

Within the scope of the present study, we did not find co-learning to facilitate learning success as compared to individual learning. Other studies which deployed IntelliCages to test spatial learning found that co-learning paradigms could even rescue the performance of amyloid precursor protein-mutants (APP.V717I), which manifested a form of Alzheimer’s disease (41). However, the task presented to the mice in the APP study was simpler and the experimental design less complicated than the method deployed in the present investigation – for instance, all animals housed together had the exact same learning goals in the Kiryk *et al.* study (41).

In the present study, the experimental design (including the limited number of operant corners per cage; the frequent changes of reward probabilities; as well as the highly dynamic environment of social groups housing both individually- and co-learning animals) may in fact have favored heuristic learning strategies among the animals (i.e., frequenting corners until a reward was received over orienting themselves based on their teammates). We did however observe a slight, non-significant trend of co-learning animals requiring less visits than solo-learning animals to collect the first 10 rewards of a trialing phase. Hence, we argue future studies with a slightly refined probabilistic place learning paradigm may in fact find co-learning to facilitate learning success over individual learning – given the feasibility of ‘trial & error’-strategies was decreased in such studies, for instance by increasing the ‘cost’ of a trial over merely restricting the time window for maximizing rewards.

While the results of the present study may seem at odds with the report of Kyrik *et al.* 2011 (41), newer reports appear to corroborate our observations of mice not performing better in co-learning scenarios over individual learning. Vezyrakis *et al*. 2025 (42) observed mice to solve puzzles more successfully (60 % solved) when being presented with them individually in an arena than when attempting to solve them in a semi-naturalistic group setting (21.4 % solved). Similar effects have also been observed in other mammalian species (43), and are hypothesized by the authors to be caused by social environments distracting the animals from the tasks at hand, as well as the animals having less relative time and energy to spend on problem solving tasks due to their involvement in complex social interactions including competition for space and potential mates (42,44,45). Of note, we did also observe a monotonic association between learning performance and several socio-positive behaviors only in the animals with the lowest overall preference for the best reward option – however, there was no overall negative association between socio-positive interactions and learning performance. Yet, the highly dynamic, ‘noisy’ social environment deployed in the present study may have manifested in co-learning animals not being able to utilize the social cues available to them to their advantage.

How exactly social behaviors relate to learning in social interactions in mice remains elusive. It has been demonstrated that mice can, to a degree, learn by observation (16,17) and are influenced in their food choices by olfactory cues derived from conspecifics (reviewed in (19)). Modes of observational learning as well as olfactory cues were also available to the team-learning animals in the present study – yet, their preference for the best reward opportunities did on average not exceed those of the individually learning animals. However, we did in fact observe ‘teammates’ to associate more often with each other in regard to some prosocial behaviors (approach rearing animals, oral-genital to oral-oral contact sequences, stopping while in contact). At this point, we cannot offer a definitive explanation whether and how these behaviors may have contributed to social learning, while it is conceivable the increased rate of oral-genital to oral-oral contact transitions may be associated with the animals smelling the sucrose reward on animals who just successfully received a reward. Yet, as stated earlier, this did not result in overall higher learning success. Further, the increased interaction rate in co-learning animals did not span all probed behaviors, and no direct overall link between social interaction rates and learning performance could be established. These results are in line with the findings of Vezyrakis *et al*. (2025) (42) and suggest learning in social interactions in mice to be a complex, multifactorial process, which may likely not be fully understood through ‘classical’ temporally- and spatially confined tests of dyadic interactions (19). We anticipate modelling of social interactions in conjunction with homecage-based approaches combining RFID- and video tracking to shed more light on social learning on mice in the near future (46).

Once current limitations of the present method are addressed, we believe to have a powerful tool at our hands to investigate social interactions and learning in rodents applicable to manifold research questions. We will further our efforts to elucidate the effects of co-learning on learning strategies in social mouse groups and aim to more conclusively identify social behaviors involved in social transmission of information. In future studies, we plan to better understand social learning strategies by deploying reinforcement learning model approaches of mouse behavior in social groups. Predictions from such models can be easily tested in the one-armed bandit-inspired probabilistic experimental design demanding a trade-off between exploration and exploitation of the best known option.

Apart from basic research on social learning in the mouse model, we anticipate a wide array of possible applications on the described method. It may for instance be deployed for investigating genes associated with the onset of autism in animal models (22,47,48) in more realistic group settings or for screening novel compounds for adverse effects on cognition and social capabilities which may only become apparent in prolonged group interactions. Overall, we will continue our efforts to further optimize this promising approach towards the fully automated investigation of learning in social interactions in mice.

## Acknowledgments

We thank the animal caretakers for their diligent care of the animals. Further, we thank Marcus Boon and Henning Sprekeler for their valuable input in developing the experimental paradigm, Pia Kahnau for insight into her experience with the TSE IntelliCage as well as Birk Urmersbach for technical assistance.

## 6 Supporting Information

**S. Fig. 1**: Exemplary detection rates during first trialing session of a night (21:00 – 23:00)

**S. Fig. 2**: Average dyadic interaction rates per confirmed copresence frame during baseline measurements. Animals were not assigned to co-learning or individual learning configuration yet, all animals received individual tasks. Color-coding is retroactively projected from the later assignment to allow for direct comparison of the animal groups. Boxplots depict average dyadic interaction type. To minimize ID inaccuracies, only tracks were considered that 1) contained at least 1 successful RFID read and 2) were within the maximum speed limit (100 cm/s) as well as 3) above the movement threshold (1 px/f). These criteria had to be met by all interaction partners in order for the event to be considered. Lines represent median; lower hinges represent 0.25 quantiles; upper hinges represent 0.75 quantiles; whiskers represent ≤ Q3 + 1.5 × IQR and ≥ Q1 − 1.5 × IQR, respectively. Y-axes represent the average interaction rates per dyad: the total inter-action durations in frames for all event types were calculated per dyad; as well as the confirmed copresence in frames for all dyads. The ratio of total interaction duration / confirmed copresence per dyad represents the interaction rate. (A) Overview plot. Along the x-axis, the observed dyadic interactions were plotted on a uniform linear y-axis. (B-S) Individual plots per interaction type with individual y-scales for comparison of the social learning configuration classes: (B) Approach (C) Approach contact (D) Approach rear (E) Social approach (F) Social escape (G) Contact (H) Move in contact (I) Stop in contact (J) Group of 2 (K) Break contact (L) Oral-oral contact (M) Oral-genital contact (N) Oral-oral to oral-genital contact sequence (O) Oral-genital to oral-oral contact sequence (P) Long chase (Q) Side-by-side contact (R) Side-by-side contact in opposite orientation (S) Get away.

